# Optical imaging of single protein size, charge, mobility, binding and conformational change

**DOI:** 10.1101/505404

**Authors:** Guanzhong Ma, Hao Zhu, Zijian Wan, Yunze Yang, Shaopeng Wang, Nongjian Tao

## Abstract

Protein analysis has relied on electrophoresis, mass spectroscopy and immunoassay, which separate, detect and identify proteins based on the size, charge, mobility and binding to antibodies. However, measuring these quantities at the single molecule level has not been possible. We tether a protein to a surface with a flexible polymer, drive the protein into mechanical oscillation with an alternating electric field, and image the protein oscillation with a near field imaging method, from which we determine the size, charge, mobility of the protein. We also measure binding of antibodies to single proteins and ligand binding-induced conformational changes in single proteins. This work provides new capabilities for protein analysis and disease biomarker detection at the single molecule level.

---

Proteins play a central role in nearly every aspect of cellular functions.^1–3^ They also serve as drugs, drug targets and disease biomarkers.^4, 5^ Detecting and identifying proteins are thus the basic tasks in biomedical research, and in disease diagnosis and therapeutics.^6–8^ Various technologies have been developed for protein analysis, and the most important ones include liquid chromatography (LC), mass spectrometry (MS) and the Western Blot.^9–13^ These technologies separate proteins based on their physical characteristics, such as charge and size, and identify them based on the mass or binding to antibodies. Although ubiquitous in both biomedical industry and research labs, they are time consuming and destructive, involving protein fragmentation and denaturation.^9, 10^ They also lack single molecule detection capability. Here we report a method to image single proteins without labels, measure the size, charge and mobility of each protein simultaneously, and analyze antibody binding to the proteins in real time. The proteins are resolved individually in space on a surface, thus requiring no separation. The simultaneous charge and size quantification, together with specific antibody binding, allow identification of the protein. The method is analogous to the LC, MC and Western Blot technologies, but achieved at the single molecule level. We further show that the method allows detection of conformational changes of single proteins.

Several technologies have been demonstrated to detect single proteins without using fluorescent labels.^14–16^ One is to detect refractive index changes of proteins resulted from local heating by light.^14^ A more direct method is to measure protein binding to plasmonic hotspots on the nanorod surface from plasmonic absorption.^15^ Because the plasmonic field is non-uniform on the surface, the protein binding-induced plasmonic absorption depends on not only the size of the protein, but also where the protein binds, which makes it difficult to quantify the size of the protein. Recently, a light interference method has been developed to quantify the protein size based on optical scattering intensity.^16^ These label-free methods are attractive for protein analysis because they measure the size, an intrinsic property of proteins. However, size alone provides only limited information. Different proteins may have a similar size, but drastically different conformations, charges and binding affinities to other proteins.^17–19^ This is the reason that the popular protein analysis technologies separate proteins based on the size (mass) and charge (e.g., Western Blot, LC and MS), and identify proteins based on their specific bindings to antibodies (e.g., Western Blot and ELISA). The method in the present work can image the size and charge of each individual protein simultaneously, and measures conformation changes in the protein and specific binding to its antibody.

To achieve single protein imaging capability without labels, we tether single proteins to an indium tin oxide (ITO) coated glass slide via a flexible polymer linker (polyenthylene glycol, or PEG) and drive the proteins into oscillation by applying an alternating electric field to the ITO surface (Figures 1a-b). The ITO slide is placed on the objective of an inverted optical microscope, and incident light is directed onto the ITO surface via the objective from an appropriate angle to generate an evanescent field near the ITO surface (Figure 1a). The evanescent field interacts with the oscillating protein and leads to scattered light, which is collected by the same objective and imaged by a CMOS imager. Because the evanescent field is localized near the ITO surface, the scattered light is extremely sensitive to the protein-surface distance. As the protein oscillates, so does the scattered light, which is recorded as an image sequence (Figure 1c). We perform Fast Fourier Transform (FFT) on each pixel of the recorded image sequence to remove noise at frequencies other than the frequency of the applied field. The FFT image resolves a single protein as a bright spot with a parabolic tail that arises from the interference between the scattering of the evanescent wave by the protein and reflection from the surface (Figure 1d) (see Supplementary Information for imaging principle) ^20^. The FFT image contrast image measures the oscillation amplitude (referred to as oscillation amplitude image), which provides size, charge, and mobility of the protein as we show below.

**Figure 1.**
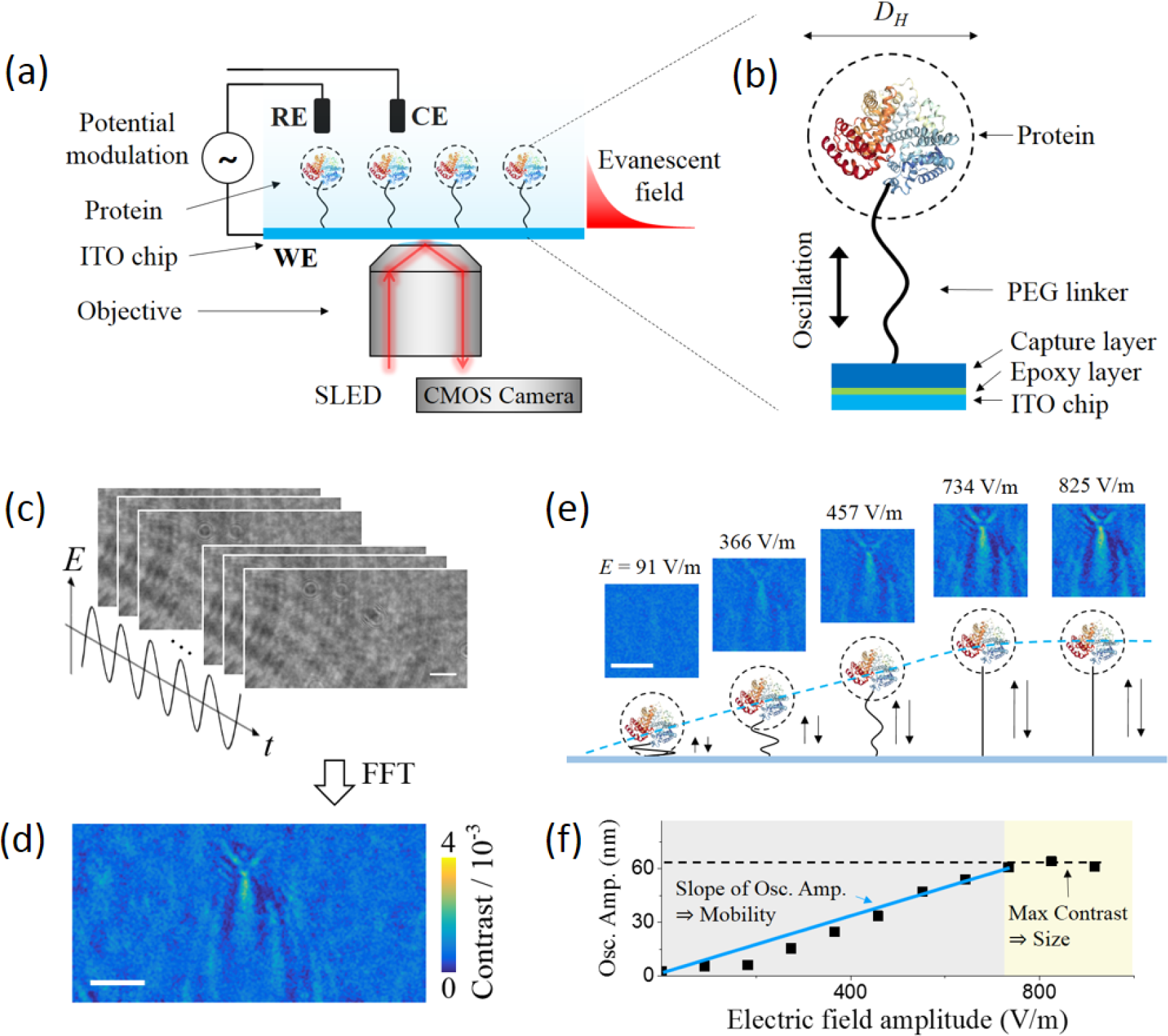
Imaging single proteins and mechanical oscillations. (a) Protein molecules are tethered to an ITO surface with a flexible polymer linker. An alternating electric field (or potential) is applied with a three-electrode electrochemical configuration to drive the molecules into oscillation, where WE, RE, and CE are the working (the ITO surface), quasi-reference (Ag wire) and counter electrode (Pt coil), respectively. The oscillating molecules scatter an evanescent field generated by illuminating the ITO surface at an appropriate angle, which are imaged with a CMOS imager. (b) The polymer linker is a 63 nm long polyethylene glycol (PEG), which couples the proteins to the ITO surface via surface chemistry described in the Method. (c) The oscillation of the individual molecules (bovine serum albumin (BSA)) is imaged at 800 frames/s, where the potential and frequency are 8 V and 80 Hz, respectively. (d) Fast Fourier transform (FFT) filter is applied to the time sequence of images shown in d to produce an oscillation amplitude image, which resolves single BSA molecules. (e) Oscillation amplitude image contrast vs. applied potential, showing an increase regime at low fields, and a plateau regime due to fully stretching of the PEG linker at high fields. (f) Oscillation amplitude (Osc. Amp.) of a BSA molecule vs. potential, from which the hydrodynamic diameter, charge and mobility of the molecule are determined. Scale bars in (c), (d) and (e) represent 3 µm.

The protein oscillation is determined by the entropic force of the PEG linker and driving force of the applied field (Supplementary Information), and its oscillation amplitude (Δ*z*_0_) is given by

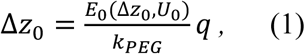

where *E*_0_(Δ*z*_0_, *U*_0_) is the amplitude of the applied field, which is a function of protein-ITO surface distance Δ*z*_0_ and surface potential *U*_0_, and *k_PEG_* is the entropic spring constant of the PEG linker (Supplementary Information). Eq. 1 shows that the oscillation amplitude is proportional to the electric field, but this is valid only at low fields (or at low applied potentials), where the oscillation amplitude is smaller than the PEG linker length. When the field or potential is sufficiently large, we expect that the linker become stretched and the amplitude reaches a plateau (Figure 1e). This behavior has been confirmed for all the proteins studied here, and Figures 1e and 1f show the results for bovine serum albumin (BSA) as an example.

The evanescent field decays exponentially from the ITO surface into the solution with a decay constant of *d* (a few hundred nm). Consequently, the oscillation amplitude image contrast, Δ*C*(Δ*z_0_, D_H_*), is given by (Supporting Information),

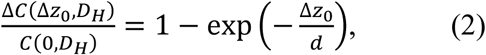

where *D_H_* is the protein hydrodynamic diameter and *C*(0, *D_H_*) is the protein FFT image contrast at zero oscillation amplitude (Δ*z*_0_ *=* 0). In the high-field plateau regime, the PEG linker is stretched, such that Δ*z* approaches the PEG length (*L_PEG_*), and the corresponding FFT image contrast, Δ*C*(Δ*z*_0_ *= L_PEG_, D_H_*), is maximum. From the measured Δ*C*(Δ*z*_0_ *= L_PEG_, D_H_*), Eq. 2 allows determination of *C*(0, *D_H_*). Because *C*(0, *D_H_*) depends on the protein size, knowing *C*(0, *D_H_*) allows determination of *D_H_* with a calibration curve (Figures 5a-b, see also Methods). Once *C*(0, *D_H_*) and Δ*C*(Δ*z_0_, D_H_*) are known, Δ*z*_0_ can be determined with Eq. 2. The charge of protein (*q*) is obtained with Eq. 1 near the transition from the low-field linear to the high-field plateau regimes (Figure 1f). The electric field at the transition point, *E*_0_(Δ*z*_0_ = *L_PEG_, U*_0_ = *U_trans_*), is measured experimentally (Supplementary Information). The protein mobility (*μ*) is related to the effective charge (*q*) and size (*D_H_*) of the protein by *µ = q/(3π*η*D_H_*), where *η* is the buffer viscosity. This relation allows determination of *μ* from *q* and *D_H_*.

We applied the method to proteins with different sizes and charges. The first example is goat immunoglobulin G (IgG), which has a molecular weight of 150 kDa and is negatively charged in the buffer (pH = 7.4). Figure 2a shows the oscillation amplitude image of several IgG molecules at *U*_0_ = 8 V. The image contrast and the extracted oscillation amplitude of IgG increase with the electric field below 8 V, and reach plateau values above 8 V (Figure 2b). From the transition points of the oscillation amplitude vs. potential plots, we obtained the charge of the individual IgG molecules. From the plateau regime, we determined the diameter of IgG, and then mobility of each IgG molecule. The oscillation amplitude is in phase (∼0^°^ phase shift) with the applied potential (Figure S2a), confirming negative charge of IgG.

**Figure 2.**
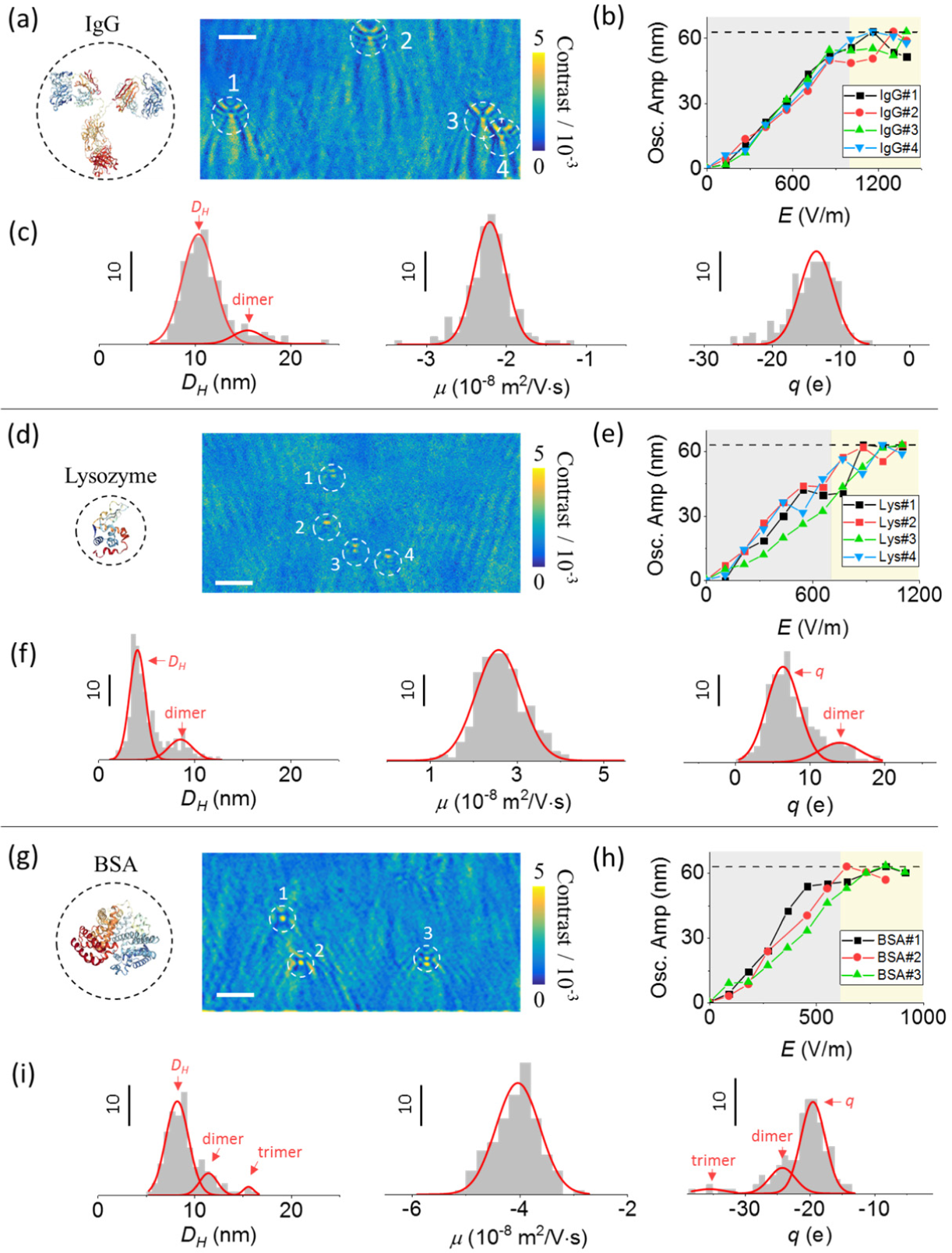
Quantifying the size, charge and mobility of single protein molecules. (a) Oscillation amplitude image of immunoglobulin G (IgG) molecules measured at *U*_0_ = 8 V. (b) Oscillation amplitude vs. applied potential plots of the IgG molecules marked in (a), from which diameter (*D_H_*), charge (*q*), and mobility (*µ*) are obtained (see Table 1). (c) Statistical analysis of *D_H_*, *q*, and *µ* measured for 186 IgG molecules, where the red curves are Gaussian fittings to the histograms (see Table 2). (d) Oscillation amplitude image of lysozyme molecules measured at potential of 9 V. (e) Oscillation amplitude vs. applied potential plots of the lysozyme molecules marked in (d), where the extracted *D_H_*, *q*, and *µ* of the molecules are listed in Table 1. (f) Statistical analysis of 246 lysozyme molecules, where the red curves are Gaussian fittings to the histograms (see Table 2). (g) Oscillation amplitude image of BSA molecules obtained at potential of 8 V. (h) Oscillation amplitude vs. applied potential plots of the BSA molecules marked in (g), where the extracted *D_H_*, *q*, and *µ* are listed in Table 1. (i) Statistical analysis of 144 BSA molecules, where the red curves are Gaussian fittings to the histograms (see Table 2). In the diameter and charge histograms, small secondary peaks are observed in these proteins, which are due to dimers. Scale bars in (a), (d) and (g) represent 3 µm.

**Table 1.** Size (*D_H_*), charge (*q*), and mobility (*µ*) of the individual protein molecules shown in Figure 2.

|  | IgG |  |  |  | Lysozyme |  |  |  | BSA |  |  |
| --- | --- | --- | --- | --- | --- | --- | --- | --- | --- | --- | --- |
|  | #1 | #2 | #3 | #4 | #1 | #2 | #3 | #4 | #1 | #2 | #3 |
| $D_H$ (nm) | 11.8 | 10.7 | 11.7 | 11.3 | 3.9 | 5.2 | 5.0 | 5.0 | 8.4 | 7.6 | 8.7 |
| $q$ (e) | -6.3 | -5.4 | -6.4 | -6.2 | 4.9 | 5.8 | 4.3 | 5.8 | -6.6 | -6.7 | -5.4 |
| $\mu$ ($10^{-8}$ m<sup>2</sup>/V·s) | -1.0 | -0.97 | -1.0 | -1.1 | 2.4 | 2.1 | 1.6 | 2.2 | -1.5 | -1.7 | -1.2 |

**Table 2.** Measured size (*D_H_*), charge (*q*), and mobility (*µ*) of protein molecules and ligand-protein complexes.

| | Molecules studied | $D_H$ (nm) | $q$ (e) | $\mu$ ($10^{-8}$ m<sup>2</sup>/V·s) |
| --- | --- | --- | --- | --- |
| IgG | 186 | $10.4 \pm 3.4$ | $-5.0 \pm 1.2$ | $-0.86 \pm 0.39$ |
| BSA | 144 | $8.3 \pm 2.4$ | $-5.3 \pm 2.0$ | $-1.2 \pm 0.75$ |
| Lysozyme | 246 | $4.1 \pm 1.7$ | $4.3 \pm 1.2$ | $1.8 \pm 1.6$ |
| CaM | 150 | $5.3 \pm 2.8$ | $-6.5 \pm 1.9$ | $-2.0 \pm 1.2$ |
| Ca<sup>2+</sup>/CaM | 151 | $6.0 \pm 2.9$ | $-5.1 \pm 1.7$ | $-1.4 \pm 1.1$ |
| Anti-IgG/IgG | 137 | $13.2 \pm 1.8$ | $-7.2 \pm 2.0$ | $-0.90 \pm 0.62$ |

Using this procedure, we analyzed 186 IgG molecules. Figure 2c plots the histograms for the diameter, charge and mobility, showing pronounced peaks at 10.4 nm, −5.0 e (e, the elementary charge, is 1.6×10^−19^ C) and −0.86×10^−8^ m^2^V^−1^s^−1^, respectively. The mean size and mobility agree with the values from dynamic light scattering experiments for IgG (Figure 5c) and reported in literature (Table S1 and S2), and the mean charge is also close to the estimated value (Table S3). The agreements of the diameter, charge and mobility with the reference experiments and literature support that the oscillation amplitude images are primarily due to single molecules. The standard deviations of the diameter (3.4 nm) and charge (1.2 e) histograms are much smaller than the mean values (10.4 nm and −5.0 e). The diameter histogram displays a small secondary peak located at a larger diameter, which is attributed to formation of dimers (Figure 2c). Small secondary peaks also appear in the diameter and charge histograms of other proteins (Figures 2f and i), which further confirm that the images are primarily due to single molecules. This conclusion is supported by the calibration plot generated using polystyrene nanoparticles of difference sizes (Figures 5a-b and see details later).

To ensure that the individual patterns shown in Figure 2a are indeed single IgG molecules, we studied anti-IgG binding to the IgG tethered on the surface (Figure 3a). We first flew PBS buffer over the IgG molecules (oscillating in the plateau regime). After establishing a baseline, we then introduced anti-IgG and monitored its binding to the IgG. Upon the introduction of anti-IgG, the apparent diameter of IgG increases (Figure 3b), indicating binding of anti-IgG to the IgG and formation of an anti-IgG/IgG complex. After measuring the binding process, we flew buffer over the surface and observed diameter decrease in some anti-IgG/IgG binding complexes, indicating unbinding of anti-IgG. The binding and unbinding events are also shown in the oscillation amplitude images captured during the measurement (Figure 3c). To confirm the observation, we performed end-point measurement by incubating IgG with 33 nM anti-IgG. The diameter histogram shows two peaks located at 10.3 nm, and 13.2 nm, respectively (Figure 3d). The former is IgG, and the later corresponds to IgG/anti-IgG. The charge histogram also reveals two peaks, located at −4.8 e and −7.2 e, which are associated with IgG and IgG/anti-IgG complex (Figure 3d). In contrast, the mobility shows only one peak (Figure 3d). This is because that mobility is intensive quantity and scales with *q*/*D_H_*. Compared to Figure 2c, the appearance of the IgG/anti-IgG peak in diameter and charge histograms verifies the binding of anti-goat IgG to goat IgG. The IgG peak (red) indicates some goat IgG molecules remain unbound after incubation, which could be due to the unfavorable orientation of the molecules as tethered by the PEG linker. To further ensure specific binding of anti-IgG to IgG, we performed a control experiment by introducing anti-human IgG and observed no changes in the size and oscillation amplitude image of the goat IgG (Figures 3e-f).

**Figure 3.**
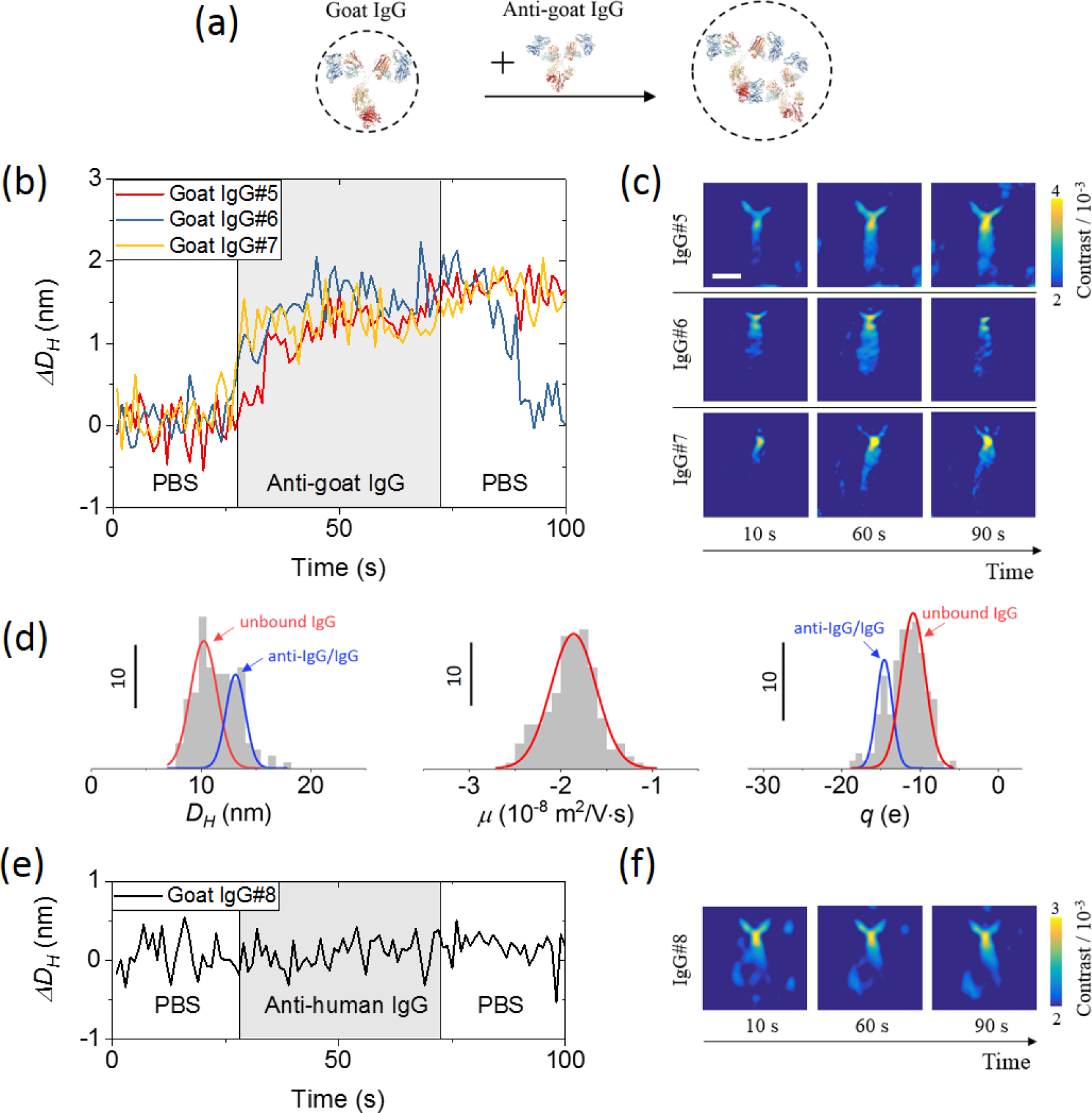
Identifying single proteins via antibody binding. (a) Anti-goat IgG is introduced to bind with PEG tethered goat IgG. (b) Binding/unbinding of anti-goat IgG with three goat IgG molecules tracked in real-time, showing diameter changes associated with the binding and unbinding events. (c) Snapshots of the three IgG molecules captured before, during and after the binding experiment in (b). The scale bar represents 3 µm. (d) Statistical analysis of 137 goat IgG molecules showing the diameter (*D_H_*), charge (*q*) and mobility (*µ*) histograms of the molecules after incubation the molecules with 33 nM anti-goat IgG for ∼30 min, the two peaks in the diameter and charge histograms correspond to IgG and anti-IgG/IgG complex. The mobility histogram has one broad peak only because mobility is an intensive quantity and related to the ratio of the charge to the diameter. The peaks are fitted to Gaussian distribution and the results are shown in Table 2. (e) A control experiment using anti-human IgG, showing no detectable changes in the diameter of IgG. (e) Snapshots of IgG#8 molecule during the binding experiment in (e).

We applied the method to lysozyme (MW=14 kDa), a much smaller protein than IgG. Lysozyme has lower image contrast than IgG because of its smaller size (Figure 2d). The image intensity oscillation is out of phase (∼180^°^ phase shift) with the applied potential (Figure S2c). This is the opposite of IgG, but expected because lysozyme is positively charged at pH = 7.4. Similar to IgG, the lysozyme oscillation amplitude increases with the field (< 9 V) and then approaches a plateau as the PEG linker reaches its maximum stretching length (Figure 2e). We determined *D_H_*, *q* and *µ* of the individual lysozyme molecules and constructed histograms for these quantities (Figure 2f). The mean values of *D_H_*, *q* and *µ* are 4.1 nm, 4.3 e and 1.8×10^−8^m^2^V^−1^s^−1^, respectively. The measured *D_H_* and *µ* are consistent with the dynamic light scattering data (Figure 5c), and the charge agrees with the expected value (Table S3).

Another example is BSA (MW=66 kDa), which is smaller than IgG but larger than lysozyme. As shown in Figure 2g, BSA has image contrast lower than IgG but greater than lysozyme, which is consistent with the size of the molecule. We plotted BSA oscillation amplitude vs. potential and observed similar dependence as IgG and lysosome: a low-field increasing regime followed by a high-field plateau regime (Figure 3h). The measured *D_H_*, *q* and *µ* are 8.3 nm, −5.3 e and −1.2×10^−8^ m^2^V^−1^s^−1^ for BSA (Figure 3i). These results agree with the values from the dynamic light scattering (Figure 5c) and calculated charge (Table S3). We summarize the results for IgG, lysozyme and BSA, as well as other proteins and complexes in Table 2.

In addition to quantifying the size, charge and mobility of single proteins, the present imaging technology can measure conformation changes in proteins. To demonstrate this capability, we studied Ca^2+^ binding to calmodulin (CaM), a protein that mediates various important Ca^2+^ signaling processes, such as muscle contraction, inflammation and fertilization.^21^ CaM has two globular domains, each containing two EF-hand motifs, so it can bind up to four Ca^2+^ and causes a conformal change in CaM (Figure 4a).^22^ We tethered CaM to an ITO surface, incubated it in buffers with and without Ca^2+^, and measured the oscillation vs. potential in each buffer (Figure 4b), from which we determined *D_H_*, *q* and *µ* for CaM and Ca^2+^/CaM complex. A total number of 150 CaM molecules and 151 Ca^2+^/CaM complexes were measured and the histograms are shown in Figure 4c. *D_H_* of CaM increases from 5.3 nm to 6.0 nm upon binding to Ca^2+^. This finding is also consistent with literature values,^22^ which is attributed to Ca^2+^ binding-induced conformation change in CaM. We verified this size increase by performing dynamic light scattering (Figure 5c). *q* for CaM is −6.5 e and changes to –5.1 e upon binding to Ca^2+^. *µ* for CaM is found to be –2.0 × 10^−8^ m^2^V^−1^s^−1^, which changes to –1.4 × 10^−8^ m^2^V^−1^s^−1^ after binding to Ca^2+^.

**Figure 4.**
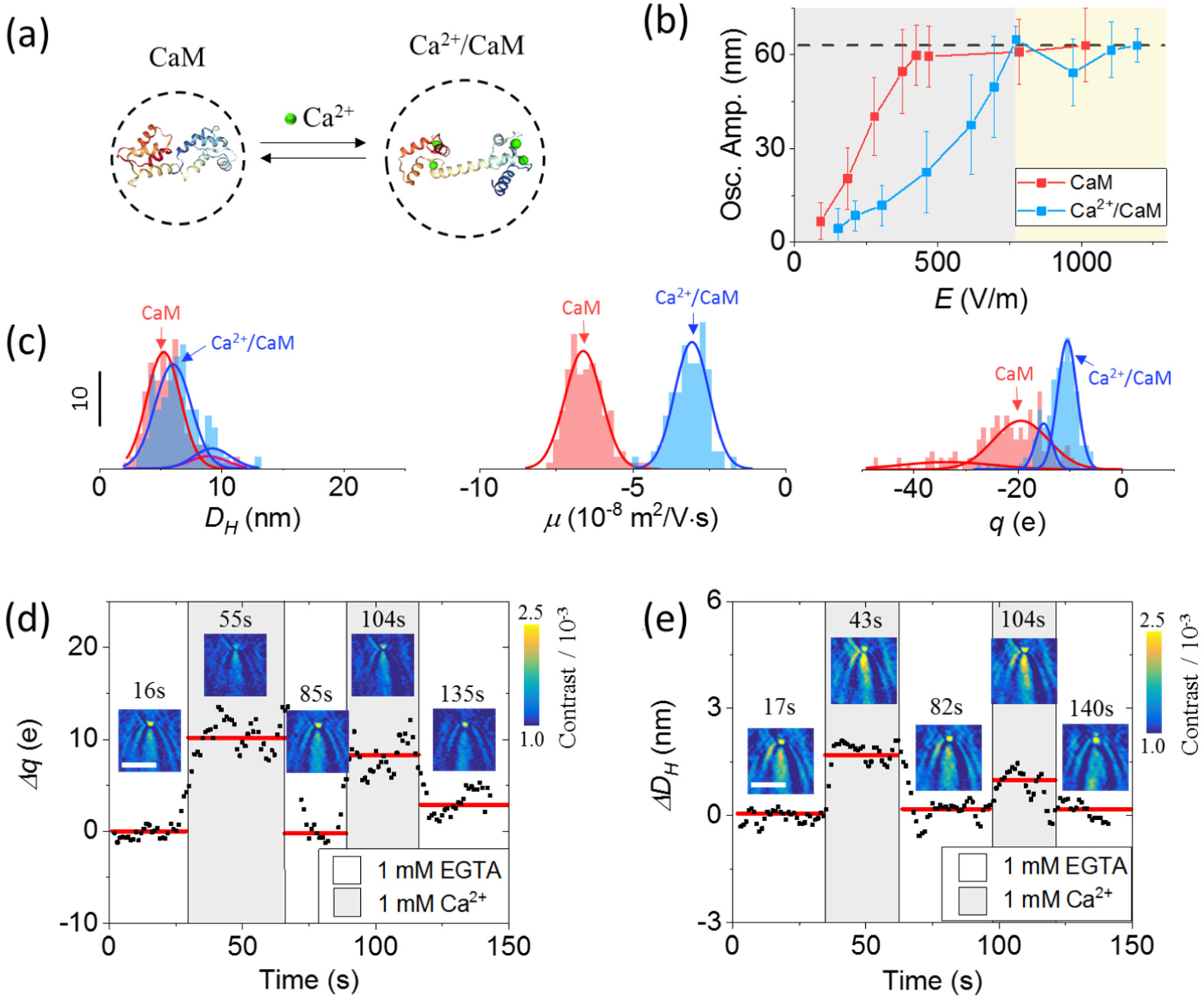
Ligand binding-induced conformation change in a protein. (a) Binding of Ca^2+^ to calmodulin (CaM) causes conformation and charge changes in CaM. (b) Oscillation amplitude vs. potential plots before (red) and after (blue) Ca^2+^ binding to CaM. The error bar represents measurement of >150 individual CaM or Ca^2+^/CaM molecules. (c) Statistical analysis for 150 CaM molecules (red) and 151 Ca^2+^/CaM molecules (blue) showing the diameter (*D_H_*), charge (*q*) and mobility (*µ*) distributions of CaM and Ca^2+^/CaM complex (see Table 2 for the summary). (d) and (e) Tracking of the charge (Δ*q*) and size (Δ*D_H_*) changes of a single CaM molecule induced by Ca^2+^ binding over time, where the potential is fixed at 4 V for the charge measurement, and at 7 V for the size measurement. For both charge and size measurements, the solution flowing over the surface is alternated between EGTA and PBS (at pH = 7.4). The scatter plot (black dots) are raw data smoothed over 3 points, and the red lines are guide to the eye, showing the charge or size change in each cycle. The inset images are snapshots of a CaM molecule captured during Ca^2+^ binding. The scale bars in (d) and (e) represent 3 µm.

We also monitored Ca^2+^ binding to CaM in real time by first driving CaM into oscillation to the maximum (plateau regime), and then alternatively flowing 1 mM Ca^2+^ and 1 mM ethylene glycol tetraacetic acid (EGTA) solutions over the surface. EGTA is known to cause unbinding of Ca^2+^ from CaM via chelation with Ca^2+^, so the experiment allowed us to repeatedly monitor the binding and unbinding processes between Ca^2+^ and a CaM molecule. From the oscillation amplitude images acquired in real time, we obtained both the effective size and charge changes of single CaM molecules (Figures 4d-e). The real-time data are consistent with the above equilibrium measurements carried out by incubating CaM in Ca^2+^ and Ca^2+^ free solutions.

We performed calibration by imaging polystyrene nanoparticles of different diameters (*D_H_* = 40-140 nm). These nanoparticles are larger than the proteins and can be directly imaged with the setup by subtracting the background from each image, allowing us to obtain the image contrasts vs. size (Figure 5a). Since the single protein measurements used a PEG linker, we evaluated the effect of the linker on the calibration plot by attaching 15-nm polystyrene nanoparticles to the ITO surface and carried out the same measurements as for the proteins (Figure 5b). The power relation between the image contrast and *D_H_* is ∼2.2, smaller than the value of 3 for a simple scattering model. This discrepancy is due to the roughness of the ITO surface as confirmed by AFM and simulation (Figure S4, Supplementary Information). The protein sizes determined with the calibration curve are close to those in literature, which validates the calibration. We further compared our results with dynamic light scattering and electrophoretic light scattering measurements (Figure 5c). The hydrodynamic diameters measured here for single proteins are in good agreement with the dynamic light scattering values and within the range reported in literature (Table S1 and S2). The single molecule mobility also agrees with those by electrophoretic light scattering for all the cases.

**Figure 5.**
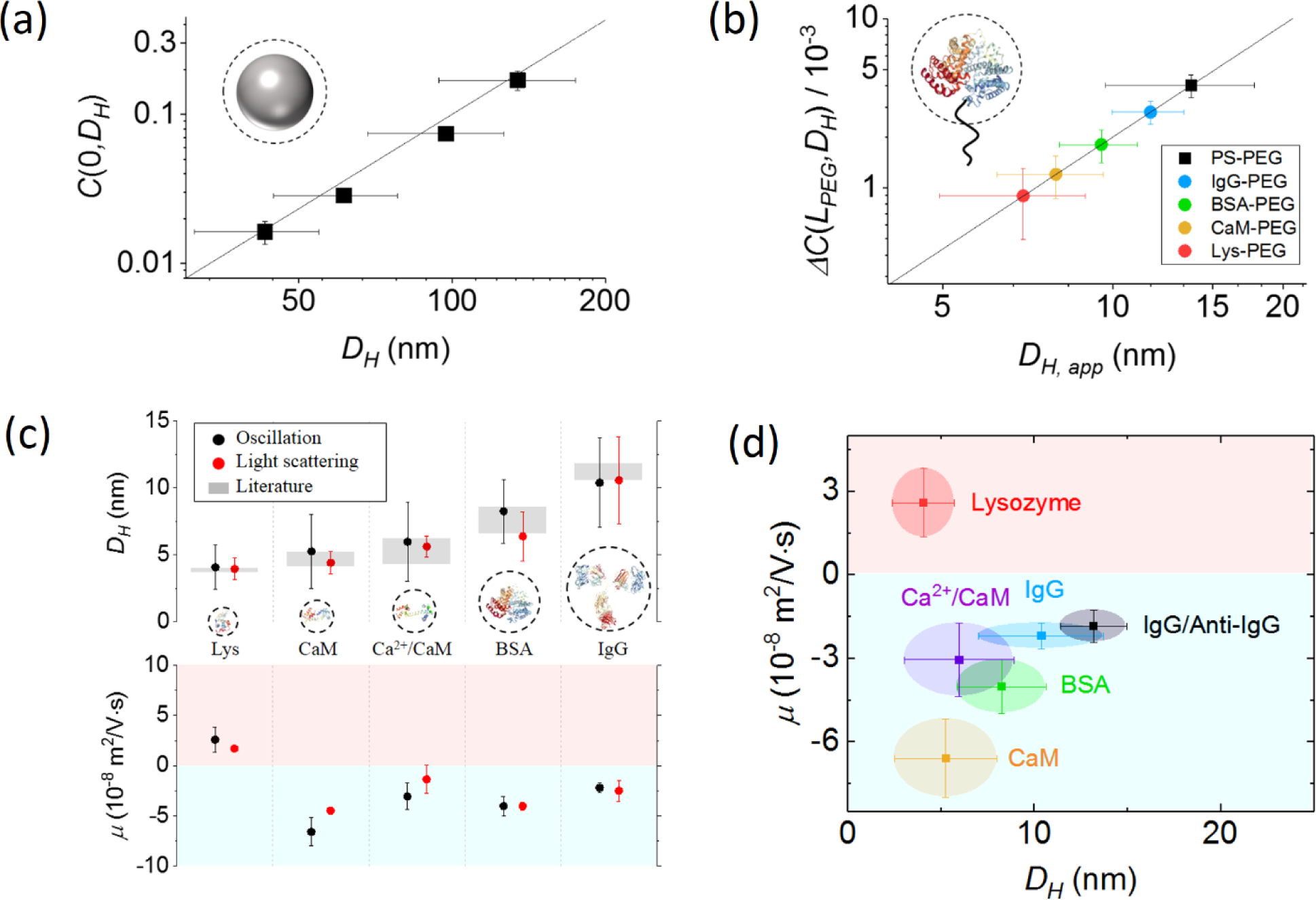
Identifying proteins based on size and mobility. (a) Image contrast vs. size for polystyrene (PS) particles. Because PS particles bind to the ITO surface from the bulk solution (Δ*z*_0_ *→ ∞*), the image contrast is *C*(0, *D_H_*) according to Eq. 2. (b) Determining protein size (*D_H,app_*) from image contrast change, *ΔC*(*L_PEG_, D_H_*). Unlike the PS particles, the proteins are tethered to the surface with a maximum distance of *L_PEG_*. We thus measured a tethered 15-nm PS particle and included the data in the plot (see Supplementary Information for details). (c) Comparison of measured *D_H_* and *µ* with light scattering experiments and also literature values. (d) Mobility (*µ*)- size (*D_H_*) plot of single proteins and protein-ligand complexes, showing different proteins or complexes are separated and thus can be identified based on the mobility and size, which resembles “2-D electrophoresis”.

Two-dimensional (2D) gel electrophoresis is a powerful technology that identifies proteins based on their size and pI (or mobility at different pH). The present single molecule imaging method can perform protein analysis in an analogous manner, but at the single molecule level and without the time-consuming separation step. This capability is shown in Figure 5d, which plots different proteins and protein-ligand complexes according to mobility and size. The proteins and complexes in the 2D-plot are separated, allowing identification of proteins like 2D electrophoresis. Binding of IgG to anti-IgG shifts the IgG region to a new position in the 2D-plot (Figure 5d). This is similar to the Western Blot and provides additional identification of the protein.

We have developed a label free technology to image single proteins, and to quantify the size, mobility and charge of single proteins simultaneously. The precisions for the size and charge achieved with the present setup are 1.0 nm and 0.3 e, respectively. The technology can also monitor protein-protein interactions and ligand binding-induced conformation changes in single proteins. Using these capabilities, we have analyzed single proteins based on size, charge, mobility and specific binding to antibodies. This resembles the widely used the Western Blot and ELISA technologies, but achieved at the single molecule level without separation and denaturation of the proteins. We anticipate that the technology will open a new revenue to study various processes of proteins, including conformation changes, molecular binding and post-translational modifications of proteins, and to detect disease biomarkers at the single molecule level without labels.

## Methods

### Materials

ITO slides with resistance of 70-100 Ω were purchased from SPI Supplies. Streptavidin was purchased from VWR. (3-Glycidyloxypropyl)trimethoxylilane, lysozyme, calmodulin, and BSA were purchased from Sigma-Aldrich. Goat IgG (anti-digoxigenin) was purchased from Abcam. Goat anti-human IgG and rabbit anti-goat IgG were purchased from Invitrogen. Polystyrene nanoparticles were purchased from Bangs Labs. Biotin-PEG-NHS (MW = 10 kDa) and streptavidin coated polystyrene particles were purchased from Nanocs. Deionized (DI) water with resistivity of 18.2 MΩ·cm was used in all the experiments.

### Experimental setup

The imaging setup was built on an inverted microscope (Olympus IX-81) with a 60x (NA=1.49) oil immersion objective. A superluminescent light emitting diode (SLED) (SLD- 260-HP-TOW-PD-670, Superlum) with central wavelength at 670 nm and output power of up to 15 mW was used as light source. A CMOS camera (ORCA-Flash 4.0, Hamamatsu) was used to record 2048 by 256 pixels images at 800 frames per second. A sinusoidal potential (*f* = 80 Hz) was applied to the ITO slide with a function generator (33521A, Agilent) and a potentiostat (AFCBP1, Pine Instrument Company) using a three-electrode configuration, where the ITO, a Ag wire and Pt coil served as the working, reference and counter electrodes, respectively. A USB data acquisition card (NI USB-6251, National Instruments) was used to synchronize the applied potential, the current, and the recorded images.

### Modification of ITO surface

The ITO slides were cleaned by sonication sequentially in acetone, ethanol, and DI water, each with 20 min, and then soaked in H_2_O_2_/NH_3_·H_2_O/H_2_O (1:3:5) for one hour, which were then rinsed with DI water and dried with N_2_. The slides were incubated in 1% (3-Glycidyloxypropyl)trimethoxylilane in isopropanol for 10 hours to silanize and form terminal epoxy groups. The epoxy-functionalized slides were rinsed with isopropanol and DI water, dried with N_2_, and incubated in 0.1 mg/ml streptavidin + 1x PBS for 4 hours. At last, the slides were incubated in 0.1 mg/ml BSA + 1x PBS for 30 minutes.

### Assembly of protein oscillators

Biotin-PEG-NHS was used to tether the protein to the functionalized ITO surface. The protein (IgG, BSA, lysozyme, or CaM) was first incubated with the biotin-PEG-NHS linker at 10:1 ratio to form a PEG-protein complex in 1x PBS overnight at 4 °C. The solution containing protein-PEG complex was then added to the streptavidin coated ITO slides and incubated for one hour to allow biotin-streptavidin binding. Finally, the chip was gently washed with 100 times diluted PBS to remove free protein molecules in the solution.

### Calibration curve

100x diluted PBS was placed on top of the ITO slide, and PS nanoparticle solution was added to allow binding of the nanoparticles to the slide surface. An image sequence was recorded at 800 frames per second for 5 seconds. The hydrodynamic diameter of each PS nanoparticle sample was measured with dynamic light scattering.

### Signal processing

FFT filter was applied to the recorded image sequence. A region of interest (ROI) with 10 ×10 pixels was selected for each protein, and the mean intensity within the ROI (*I_p_*) was used to determine the contrast of the protein. An adjacent region of the same size was selected as a reference region, and the mean intensity of the reference region (*I_r_*) was also determined. The contrast of the protein was determined with *ΔC*(*Δz*_0_, *D_H_*) = (*I_p_* − *I_r_*) / *I*, where *I* is the mean intensity within the ROI without FFT filter. The size and charge of each protein were determined based on the contrast (Supplementary Information).

