## Supporting Information for "Optical imaging of single protein size, charge, mobility, binding and conformational change"

**Entropy of PEG linker and oscillation of tethered protein molecules**

The oscillation of a protein molecule tethered by a PEG linker can be described by,

$$m\frac{d^2z}{dt^2} + c\frac{dz}{dt} + k_{PEG}z = qE, \quad \text{(S1)}$$

where $m$, $z$, $c$, $k_{PEG}$, $q$, and $E$ are mass, displacement of the protein molecule, damping coefficient, entropic spring constant of PEG linker, charge of the protein, and electric field, respectively. For a protein molecule with molecular weight of 100 kDa ($m = 1.7 \times 10^{-19}$g) oscillating at 80 Hz, the first term and the second term are about $10^{-13}$ pN and $10^{-3}$ pN, respectively, much smaller than the entropic force (see below).

We use the freely jointed chain (FJC) model[1, 2] to calculate the entropic force of the PEG,

$$f_{entropy} = k_{PEG}z = \frac{3k_BT}{nb^2}z, \quad \text{(S2)}$$

where $k_B$ is the Boltzmann constant, $T$ is temperature, $b$ is the Kuhn length of PEG, $n$ is the number of segments with length of $b$, and $z$ is the distance between the tethered protein molecule and surface. For PEG10k, $b$ = 0.55 nm, $n$ = 113,[3] and $k_{PEG} = 3.62 \times 10^{-4}$N/m. The entropic force is 22.8 pN when the PEG is stretched ($z$ ~ 63 nm). Thus, by ignoring the first term and the second term, Eq. S1 becomes,

$$k_{PEG}z = qE, \quad \text{(S3)}$$

Because the modulation is sinusoidal, $z = \Delta z_0 e^{j\omega t}$ and $E = E_0 e^{j\omega t}$, where the angular frequency $\omega = 2\pi f$. Also, the electric field applied to the molecule is a function of applied potential amplitude $U_0$ and molecule-surface distance $\Delta z_0$, $E = E_0(\Delta z_0, U_0)e^{j\omega t}$. By combining the above relations with Eq. S3, we obtain the oscillation equation of the protein molecule (Eq. 1).

**Measuring of electric field near the surface**

According to Eq. 1 and Figure 1f, the charge of the molecule can be obtained once the electric field ($E_0(\Delta z_0 = L_{PEG}, U_0 = U_{trans})$) at the transition point ($\Delta z_0 = L_{PEG}$) is determined. To measure the electric field, we tethered 40 nm streptavidin coated gold nanoparticles (AuNPs) to a gold film with PEG10k linkers,[4] pulled the AuNPs away from the surface by applying potential, and recorded particle-surface distance ($z$) change with the potential (Figure S1a-b). Because AuNPs are negatively charged at pH = 7.4, the negative potential pulls the particles away from the surface, leading to decrease in image intensity (Figure S1c). The intensity decreases exponentially with the potential and reaches the minimum value when the PEG is fully stretched (Figure S1d). By converting the image intensity into particle-surface distance ($z$),[4] we obtained $\Delta z_0$ vs. $U_0$ plot for the AuNP (Figure S1e). The plot shows a linear regime followed by a plateau regime, which is consistent with the observation for protein molecules (Figures 2b-h and 4b). The potential at the transition point is $U_{trans}$ = -0.95 V.

To determine the field at a given potential, we calculated the charge of the AuNPs based on the zeta potential, given by,[5, 6]

$$q_{NP} = 4\pi a^2 \cdot \frac{2\varepsilon_r \varepsilon_0 \kappa k_B T}{Z} \sinh\left(\frac{Ze\zeta}{2k_B T}\right)\left[1 + \frac{1}{\kappa a \cdot \cosh^2(Ze\zeta/4k_B T)}\right], \quad \text{(S4)}$$

where $a$ is the radius of the particle, $Z$ is the valence of ions in the electrolyte solution, $\varepsilon_0$ and $\varepsilon_r$ are the permittivity of vacuum and the relative permittivity of the solution, $\kappa^{-1}$ is the Debye length, $e$ is the elementary electric charge, and $\zeta$ is zeta potential of the particle. We found that $\kappa^{-1}$ = 7.89 nm (see “charge screening effect” section) and $\zeta$ = -13.1 mV, which was measured by ELS. Thus, we obtained $q_{NP}$ with Eq. S4, which was -42.5 e. The entropic force of the stretched PEG linker is 22.8 pN, which is balanced by the electrostatic force according to $q_{NP} E_0(\Delta z_0 = L_{PEG}, U_0 =$

$U_{trans}) = k_{PEG}L_{PEG}$, from which we have $E_0 = -3.35 \times 10^6$ V/m. Because the electric field at a given distance from the surface ($\Delta z_0 = L_{PEG}$) scales with applied potential $U_0$, we obtain the following relation,

$$E_0(\Delta z_0 = L_{PEG}) = 3.53 \times 10^6\ U_0/\text{m}. \text{ (S5)}$$

Using this equation and the transition potential ($U_{trans}$) obtained with the oscillation amplitude vs. potential plot (Figure 1f), the electric field at the transition point was determined.

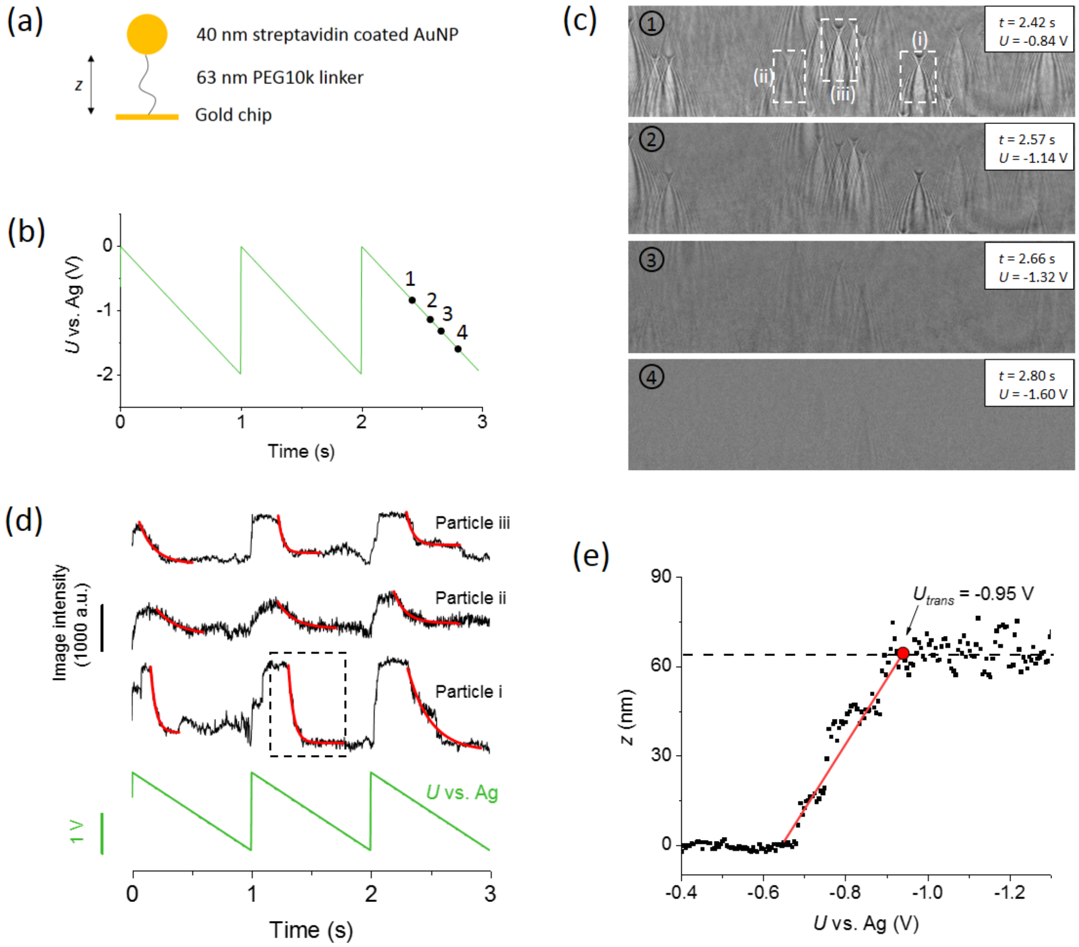


Figure S1. Measuring electric field near the surface using tethered gold nanoparticles (AuNPs). (a) 40 nm streptavidin coated AuNPs are tethered to gold chip surface via PEG linkers. Each particle is tethered by a SH-PEG10k-biotin linker (length = 63 nm) which contains a thiol group and a biotin group on its two ends. The thiol group covalently binds to the gold surface and the biotin group binds to the particle via biotin-streptavidin interaction. (b) A triangular potential sweep is applied to the gold surface to drive the AuNP. (c) Snapshots of the particles at different potentials (marked in (b)). The intensity change reflects the change in particle-surface distance. (d) Intensity response of three particles marked in (c) (black curve) to applied potential (green curve). The intensity change in each cycle is fitted to exponential decay (red curve). (e) Intensity change of particle i (marked in (d)) is converted to particle-surface distance ($z$). After the particle is pulled off the surface, the distance shows linear relationship with the applied potential and finally reaches the plateau where the PEG is fully stretched. The red line is the fitting of the linear regime, and the red dot marks the transition point, where the entropic force reaches a plateau, and the corresponding applied potential is $U_{trans}$ = -0.95 V.

**Determination of protein size and mobility**

By changing the applied electric field and performing FFT, the oscillation amplitude images of the proteins are obtained at different electric field. This allows us to plot the image contrast, $\Delta C(\Delta z_0, D_H)$ vs. the applied potential amplitude ($U_0$). From the plateau regime of the plot, corresponding to a stretched PEG linker, we determined the image contrast in the plateau regime, $\Delta C(L_{PEG}, D_H)$, which is used to determine the image contrast at $z = 0$, $C(0, D_H)$, using Eq. 2. The apparent diameter of the protein, including the contributions of the protein and the PEG linker, is obtained using the calibration curve (Figures 5a-b). Knowing the length of PEG, we extracted the protein diameter ($D_H$) with Eq. S6.

To determine the charge, $\Delta C(\Delta z_0, D_H)$ vs. $U_0$ plot is first converted into $\Delta z_0$ vs. $U_0$ plot, where $\Delta z_0$ is obtained using Eq. 2, according to,

$$\frac{1-\exp\left(-\frac{\Delta z_0}{d}\right)}{1-\exp\left(-\frac{L_{PEG}}{d}\right)} = \frac{\Delta C(\Delta z_0, D_H)}{\Delta C(L_{PEG}, D_H)}. \quad \text{(S4)}$$

The transition point of the $\Delta z_0$ vs. $U_0$ plot is determined from the intersection between linear regime and the plateau regimes.

**Effect of PEG linkers**

The size obtained from the oscillation image contrast include contributions from the linker. To extract the diameter of the protein ($D_H$), the following equation is used,

$$D_H^3 = D_{H,app}^3 - D_{H,PEG}^3, \quad \text{(S6)}$$

where $D_{H,app}$ is the apparent diameter of the protein, and $D_{H,PEG}$ is the diameter of PEG coil measured with DLS.

**The length of PEG linker**

The PEG monomer has a length of 0.278 nm,[7, 8] and the PEG linker used in this work has a molecular weight of 10 kDa, consisting of 225 ethylene glycol units. The linear length of the PEG is: 0.278 nm × 225 = 63 nm.

**Extracting oscillation amplitude by performing FFT**

The images captured by the CMOS imager record the oscillation of protein molecules over time. Plotting the local image contrast vs. time reveals periodic oscillation of an IgG molecule (red dashed line, Figures S2a and S2c). The FFT amplitude spectrum shows a sharp peak located at the frequency of the applied electric field (red line, Figures S2b and S2d). We performed this FFT analysis on each pixel of the time sequence images (Figure 1c), extracted the oscillation amplitude averaged over one second, and constructed an FFT image (oscillation amplitude image) shown in Figure 1d.

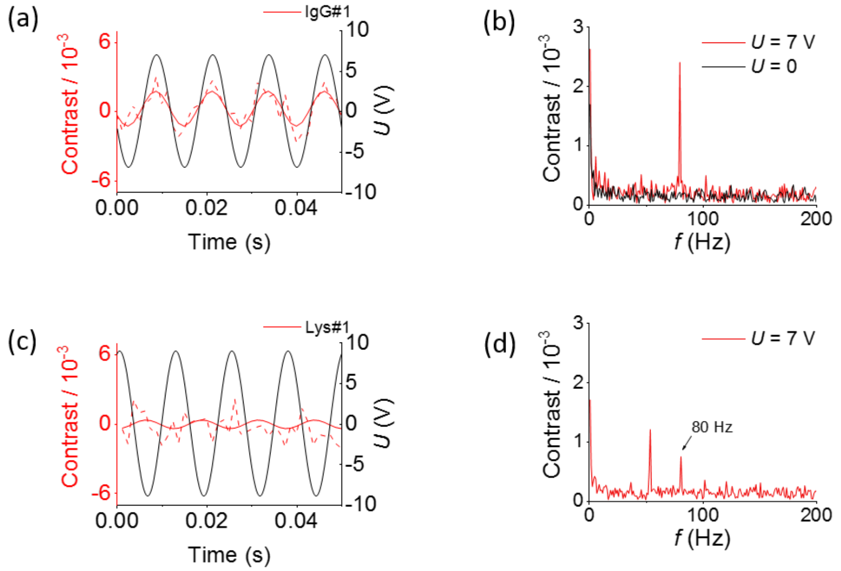


Figure S2. (a) The oscillation of an IgG molecule with potential (*U*). The dashed red line shows the oscillation of the molecule and the solid red line is the component at 80 Hz obtained by applying FFT filter. The phase difference between contrast and *U* is ~ 0°, which indicates IgG is negatively charged. (b) FFT of the oscillation in (a) ***over one second*** shows a pronounced peak at 80 Hz (red curve). No peak at 80 Hz is observed without applying electric field (black curve). (c) The oscillation of a lysozyme (Lys) molecule with potential (*U*). The dashed red line shows the oscillation of the molecule and the solid red line is the component at 80 Hz obtained by applying FFT filter. The phase difference between contrast and *E* is ~180°, indicating that lysozyme is positively charged. (d) FFT of the oscillation of the lysozyme molecule in (c) over one second shows a pronounced peak at 80 Hz. Buffer: 100 times diluted PBS, pH = 7.4.

### Additional examples of $Ca^{2+}$ binding with CaM

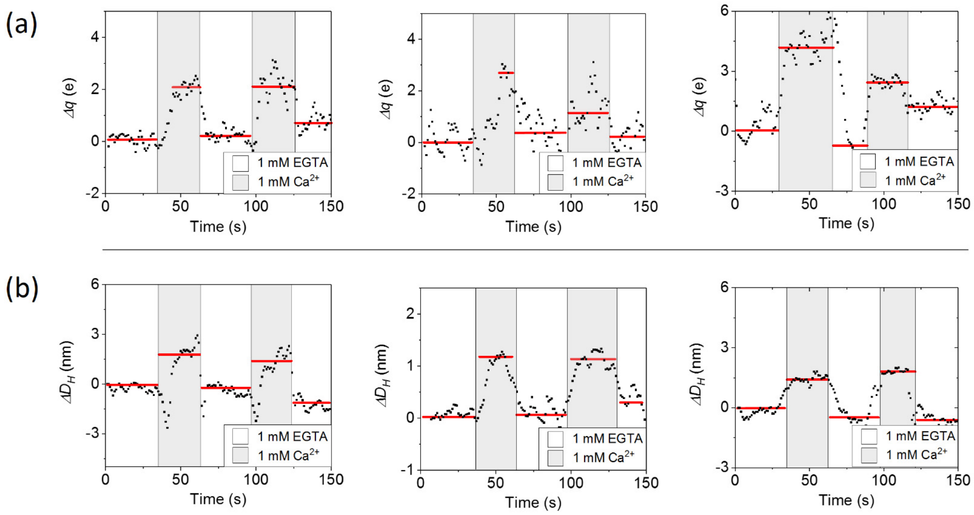


Figure S3. Additional examples showing charge change (a) and size change (b) during CaM-$Ca^{2+}$ interaction. For both charge and size measurements, the solution cycles between 1 mM EGTA in 100 times diluted PBS and 1 mM $CaCl_2$ in 100 times diluted PBS at pH = 7.4. The black points are raw data smoothed by 3 points, and the red lines are guide to the eye showing the charge or size change in each cycle.

### Imaging principle

Light is directed on the ITO surface at an angle close to the critical angle to generate an evanescent wave ($u_e$) propagating along the surface. When a protein is present on the surface, it scatters the evanescent wave, generating a scattered wave ($u_S$), given by

$$u_s(r, r') = \alpha u_e(r') e^{-\kappa|r-r'|} e^{-ik|r-r'|} \qquad \text{(S8)}$$

where $r'$ is the location of the protein, $\alpha$ is a scattering coefficient related to the polarizability of the molecule, $\kappa$ is the decaying constant of the evanescent wave, and $k$ is the wavenumber of evanescent wave. The superposition of the two waves, together with light reflected from the ITO surface ($u_r$), is

$$u(r,r') = u_r(r) + u_e(r) + u_s(r,r'). \quad \text{(S9)}$$

The overall reflected light detected by the camera ($I$) is given by,[9-13]

$$I = |u_r(r) + u_e(r) + u_s(r,r')|^2. \quad \text{(S10)}$$

The image contrast of the particle is described by

$$I(r,r') = |u_r(r) + \mathrm{u}_e(r) + u_s(r,r')|^2 - |u_r(r) + \mathrm{u}_e(r)|^2, \quad \text{(S11)}$$

where the last term is the background image in the absence of the protein, which is subtracted out. For weak scattering, $|u_s|^2$ is small, and Eq. S12 is reduced to

$$I(r,r') = 2\mathrm{Re}\{[u_r(r) + \mathrm{u}_e(r)]u_s(r,r')\}, \quad \text{(S12)}$$

which shows that the image contrast is originated from the interference between $u_r + u_e$ and $u_s$. This analysis indicates the imaging principle is interferometric, similar to iSCAT[14] and surface plasmon resonance imaging[15]. Using Eq. S12, we computed an image, which closely resembles the experimental image (Figure S4b). Eq. S12 shows the image intensity scales with $\alpha$, which is proportional to the cubic power of the diameter ($D_H^3$). The observed size dependence of the image contrast is slower than cubic power (between 2-3), which is attributed to surface roughness, as discussed below.

**Effect of surface roughness**

The above analysis assumes a perfect surface. In practice, ITO surface is rough, as shown by atomic force microscopy (AFM) (Figure S4a). The surface roughness effect is particularly important for small objects, such as protein molecules, which are comparable with or smaller than the surface rough features (grains). We simulate the surface roughness effect by including an additional term, $u_{rough}$ in Eq. S11,

$$I(r,r') = \left|u_r(r) + u_e(r) + u_{rough}(r,r') + u_s(r,r')\right|^2 - \left|u_r(r) + u_e(r) + u_{rough}(r,r')\right|^2,$$

(S13)

which leads to increased background and also slower dependence of the image contrast on the protein size (Figure S4). Using the grain size of the ITO measured from the AFM images, we performed numerical simulation of the size dependence of the image contrast. The simulation used 100 small polystyrene particles randomly distributed on the surface around a polystyrene particle of interest with diameter varying from 20 to 150 nm. The size distribution of the small particles used to simulation was based on the AFM measurement, which varied from 0-20 nm, with an average diameter of 10 nm. The logarithmic plot of the image contrast vs. diameter shows a slope of ~2.0, confirming decreased size dependence of the image contrast on the particle diameter (Figure S4c).

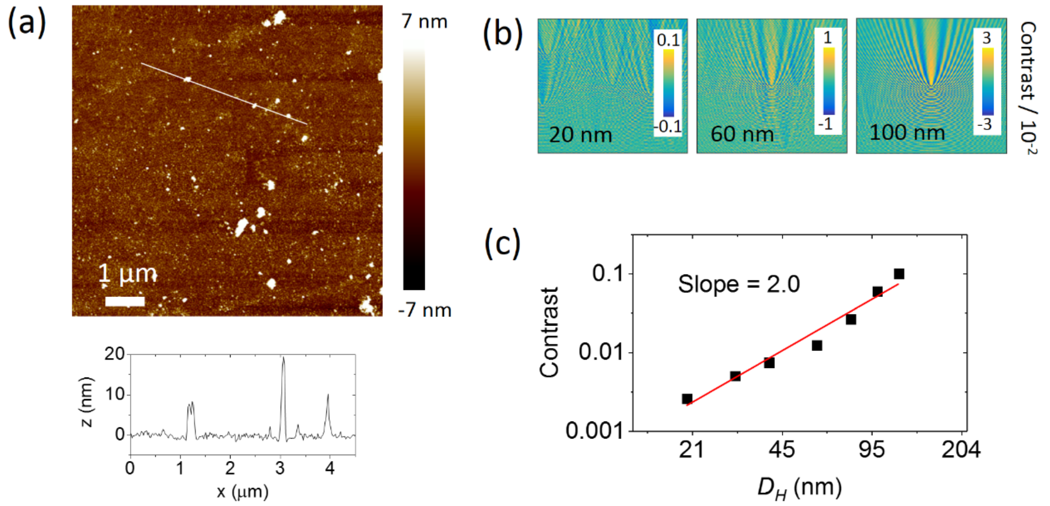


Figure S4. Surface roughness effect. (a) A typical AFM image of the ITO surface, showing grains with 0-20 nm in diameter. (b) Simulated images of polystyrene particles on the ITO surface. (c) The image contrast of polystyrene particles vs. diameter. Note that each data point represents the average over 10 simulations.

**Surface charging effect and background noise**

A bare ITO surface also responds to the applied oscillating electric field and gives rise to background noise. This response arises from the charge-dependent refractive index of ITO. To evaluate this effect, we modulated a bare ITO slide with potential ($U_0$ = 10 V and frequency = 80 Hz) and obtained the FFT images (Figure S5a). The features shown in the images are due to the grains of the ITO surface, which affect limits of detection for the diameter ($D_H$) and charge ($q$). We converted the features in the ITO background image to the equivalent $D_H$ and $q$ noise images using the calibration curve in Figure 5b and the Einstein equation (assume the mobility is $1\times10^{-8}$ $m^2V^{-1}s^{-1}$, the typical value for proteins). The results show the distributions of $D_H$ and $q$ associated with surface roughness (Figures S5b-c).

In single molecule detection, the charge-induced features overlap with the protein images and affect detection accuracy (Figure S5g). To reduce this effect, we performed 2D FFT to the oscillation amplitude image and convert the image from real space to *k*-space, which shows two rings originated from the interference of scattered field and evanescent field (Figure S5h).[15] We apply a filter to block part of the low frequency region (Figure S5i) where the background features are located, and the result shows most of the features are removed (Figure S5j). We applied the same filter to the images in Figure S5a-c to remove the background features and obtained the distributions of $D_H$ and $q$, from which the limits of detection were estimated to be ~1.0 nm for diameter, and 0.3 e for charge (Figures S5e-f).

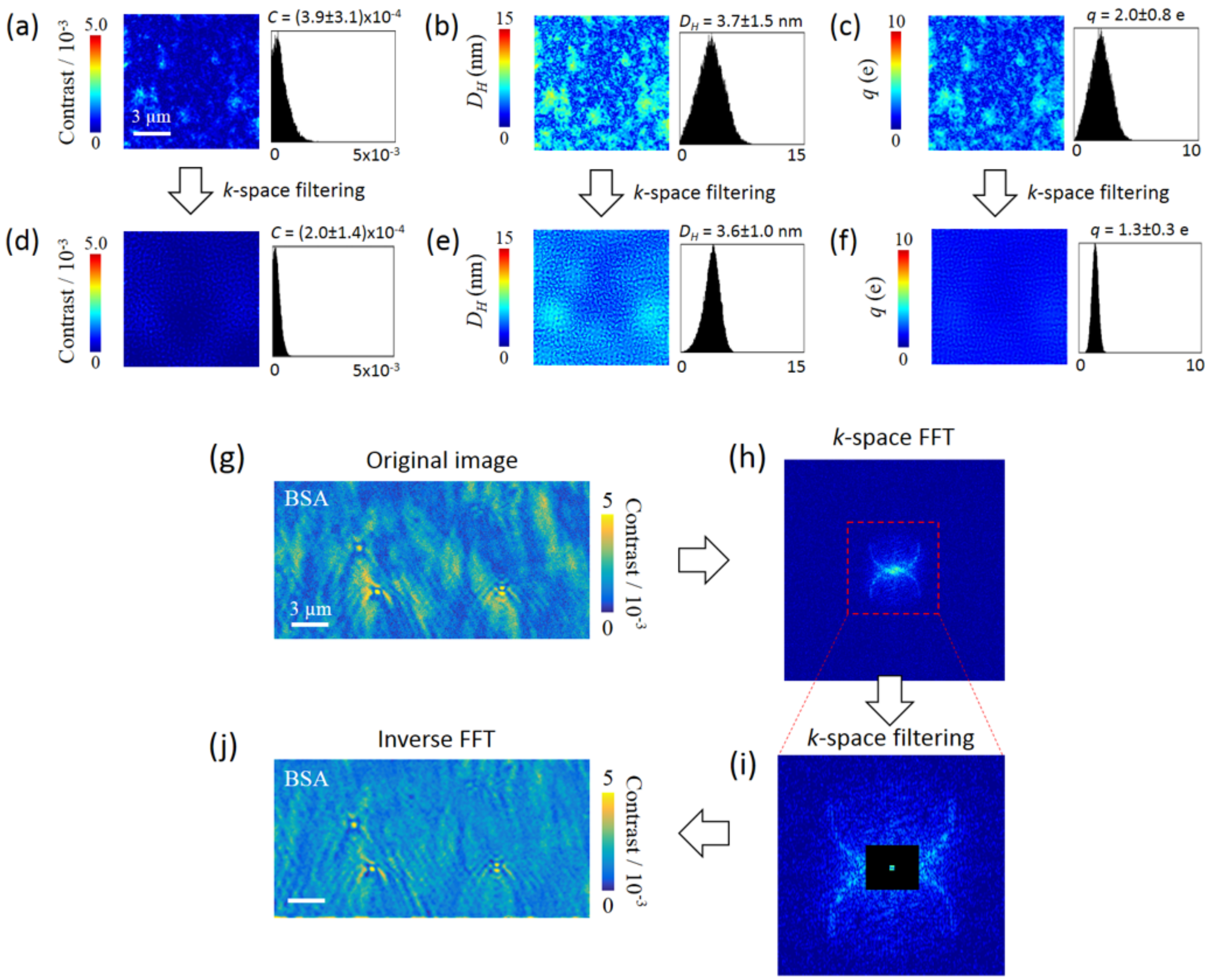


Figure S5. The effect of ITO surface charging on image contrast, and measurements of diameter and charge of proteins. (a) FFT image of a bare ITO surface modulated by applying a potential with amplitude, $U_0 = 10$ V and frequency, $f = 80$ Hz. (b) The image contrast of each pixel in (a) is converted into diameter ($D_H$), showing a histogram with $D_H = 3.7 \pm 1.5$ nm. (c) The image contrast of each pixel in (a) is converted into charge ($q$) for mobility of $1\times10^{-8}$ $m^2V^{-1}s^{-1}$. The histogram shows $q = 2.0 \pm 0.8$ e. (d)-(f) The images in (a)-(c) are filtered in $k$-space with filter shown in (i) to reduce the background features. The contrast, diameter and charge show histograms with $C = (2.0 \pm 1.4) \times 10^{-4}$, $D_H = 3.6 \pm 1.0$ nm, and $q = 1.3 \pm 0.3$ e, respectively. (g) Oscillation amplitude image of single BSA molecules and background. (h) 2D FFT is performed with the oscillation amplitude image in (g), which shows two rings originated from interference. (i) Zoom-in of the dashed region in (h). The black region shows the frequencies excluded by the filter. (j) Inverse 2D FFT of (i) shows BSA molecules and reduced background noises.

### Shot noise estimation

In the current setup, the photon collection efficiency is limited by the imager (sCMOS camera, Hamamatsu). It has a full well capacity $30000$ electrons per pixel. The recorded imaging area is 2048 × 256 pixels and the frame rate is 800 fps, which leads to a maximum photon flux of $1.2 \times 10^{13}$ e/s, corresponding to $2 \times 10^{7}$ photons per pixel per second. A typical scattering pattern of a protein or particle has an area greater than 100 × 100 pixels, which corresponds to $2 \times 10^{11}$ photons per second (recording time is 1 second). This gives an upper limit of signal-to-noise ratio of $4 \times 10^{5}$ per protein or particle. We used 10 × 10 pixels as regions of interest for contrast analysis, which has an upper limit of signal-to-noise ratio of $4 \times 10^{4}$.

### Charge screening effect

Effective charges are measured here, which are related to the net charges by,

$$\frac{\sigma_{eff}}{\sigma_{total}} = \frac{\zeta}{\psi} = e^{-\kappa x}, \qquad \text{(S14)}$$

where $\sigma_{eff}/\sigma_{total}$ is the ratio of the effective charge density to the net charge density, $\zeta$ is zeta potential of the protein, $\psi$ is the potential at the protein surface, $x$ is the slipping layer thickness, and $\kappa^{-1}$ is the Debye length, which is determined by the Debye-Hückel equation,

$$\kappa^{-1} = \sqrt{\frac{\varepsilon\varepsilon_0 k_B T}{2N_A e^2 I}}, \qquad \text{(S15)}$$

where $\varepsilon$ is the dielectric constant of the buffer, $\varepsilon_0$ is the permittivity of free space, $k_B$ is the Boltzmann constant, $N_A$ is the Avogadro number, $e$ is the elementary charge, and $I$ is the ionic strength. Using Eq. S14, the slipping layer thickness of protein molecules can be determined by the zeta potential and Debye length at two different concentrations with,

$$x = \frac{ln\zeta_1/\zeta_2}{\kappa_2 - \kappa_1}. \qquad \text{(S16)}$$

We measured the zeta potential of lysozyme in 1x PBS and 100 times diluted PBS, which are $\zeta_1 = 2.78$ mV and $\zeta_2 = 13.8$ mV, respectively. The corresponding Debye lengths are $\kappa_1^{-1} = 0.789$ nm and $\kappa_2^{-1} = 7.89$ nm according to Eq. S15, from which the slipping layer thickness for the protein is determined to be 1.4 nm. Using these parameters, we plotted $\sigma_{eff} /\sigma_{total}$ ratio vs. ionic strength (Figure S6), showing that the effective charge is ~20% of total charge for 1x PBS, and ~90% in 100 times diluted PBS.

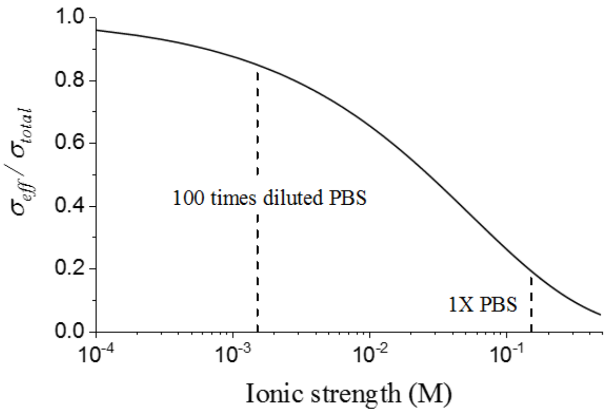


Figure S6. Charge screening effect determined with Eq. S14 using parameters listed in the text.

Table S1. Hydrodynamic diameter ($D_H$) of proteins reported in literature. *

| Protein | $D_H$ (nm) | Method | Ref. |
|---|---|---|---|
| BSA | 6.6 to 8.6 | DLS | 16 |
| | 7 | DLS | 17 |
| | 7.3 | DLS | 18 |
| IgG | 10.58 | Calculation | 19 |
| | 10.9 | DLS | 20 |
| | 10.8 to 12.5 | DLS | 21 |
| Lysozyme | 3.7 to 3.9 | DLS | 22 |
| | 3.7 | DLS | 23 |
| | 4 | Capillary electrophoresis | 24 |
| | 4 | NMR | 25 |
| CaM | 4.96±0.18 | NMR | 25 |
| | 4.4 | Calculation | 26 |
| | 4.8 | Gel permeation chromatography | 26 |
| | 5.0±0.2 | DLS | 27 |
| | 4.44 | SAXS | 28 |
| | 4.1 | SAXS | 29 |
| | 4.1 | SAXS | 30 |
| $Ca^{2+}$/CaM | 4.90±0.08 | NMR | 25 |
| | 6.0±0.2 | DLS | 27 |
| | 4.3 | SAXS | 29 |

*DLS = dynamic light scattering, SAXS = small-angle X-ray scattering, NMR = nuclear magnetic resonance.

Table S2. Mobility ($\mu$) of proteins reported in literature. *

| Protein | $\mu$ ($\times10^{-8}$ $m^2V^{-1}s^{-1}$) | Method | Ref. |
|---|---|---|---|
| BSA | -1.4 | ELS | 16 |
| | -1.7 | ELS | 31 |
| IgG | -0.8 | Capillary electrophoresis | 20 |
| | -0.32 | Capillary electrophoresis | 32 |
| Lysozyme | 0.8 | Electrophoresis and simulation | 33 |
| | 0.15 | ELS | 34 |
| | 1.8 | Capillary electrophoresis | 24 |
| | 1.8 | Capillary electrophoresis | 35 |

*Note that mobility is sensitive to pH, ionic strength and ion species, and the mobility measured in literature is not under the same experimental condition as in this work. Thus, small variation could be expected.

Table S3. Charge estimation of proteins at pH = 7.4*

| **Protein** | **Charge at pH = 7.4** | |
|---|---|---|
| | Net charge of amino acids | Estimation with zeta potential and size measured by Zetasizer** |
| **BSA** | -14.0 | -8.1 |
| **IgG** | -0.6 | -5.1 |
| **Lysozyme** | 8.2 | 2.4 |
| **CaM** | -24.1 | -6.6 |
| **CaM with 4 $Ca^{2+}$** | -16.1 | -4.1 |

*Note that the charge obtained with amino acids could be different from those measured by Zetasizer due to the binding of ions in solution.[36]

**The charge is estimated with Eq.S4 by knowing the zeta potential and size of each protein.

### *Supporting Information References*


1. Saleh, O.A. Perspective: Single polymer mechanics across the force regimes. *The Journal of chemical physics* **142**, 194902 (2015).
2. SamorÃ, P. Scanning probe microscopies beyond imaging: manipulation of molecules and nanostructures. (John Wiley & Sons, 2006).
3. Hu, Y. et al. Study of fibrinogen adsorption on poly (ethylene glycol)-modified surfaces using a quartz crystal microbalance with dissipation and a dual polarization interferometry. *Rsc Advances* **4**, 7716-7724 (2014).
4. Shan, X. et al. Detection of charges and molecules with self-assembled nano-oscillators. *Nano letters* **14**, 4151-4157 (2014).
5. Makino, K. & Ohshima, H. Electrophoretic mobility of a colloidal particle with constant surface charge density. *Langmuir* **26**, 18016-18019 (2010).
6. Loeb, A.L., Overbeek, J.T.G., Wiersema, P. & King, C. The electrical double layer around a spherical colloid particle. *Journal of The Electrochemical Society* **108**, 269C-269C (1961).
7. Hu, Y. et al. Study of fibrinogen adsorption on poly(ethylene glycol)-modified surfaces using a quartz crystal microbalance with dissipation and a dual polarization interferometry. *RSC Advances* **4**, 7716-7724 (2014).
8. Harder, P., Grunze, M., Dahint, R., Whitesides, G.M. & Laibinis, P.E. Molecular Conformation in Oligo(ethylene glycol)-Terminated Self-Assembled Monolayers on Gold and Silver Surfaces Determines Their Ability To Resist Protein Adsorption. *The Journal of Physical Chemistry B* **102**, 426-436 (1998).
9. Berger, C.E., Kooyman, R.P. & Greve, J. Surface plasmon propagation near an index step. *Opt. Commun.* **167**, 183-189 (1999).

30. Majava, V. & Kursula, P. Domain Swapping and Different Oligomeric States for the Complex Between Calmodulin and the Calmodulin-Binding Domain of Calcineurin A. *PLOS ONE* **4**, e5402 (2009).
31. Takeda, K. et al. Size and mobility of sodium dodecyl sulfate—bovine serum albumin complex as studied by dynamic light scattering and electrophoretic light scattering. *Journal of Colloid and Interface Science* **154**, 385-392 (1992).
32. Martin, N. et al. Prevention of thermally induced aggregation of IgG antibodies by noncovalent interaction with poly (acrylate) derivatives. *Biomacromolecules* **15**, 2952-2962 (2014).
33. Yamaguchi, A. & Kobayashi, M. Quantitative evaluation of shift of slipping plane and counterion binding to lysozyme by electrophoresis method. *Colloid and Polymer Science* **294**, 1019-1026 (2016).
34. Cugia, F., Monduzzi, M., Ninham, B.W. & Salis, A. Interplay of ion specificity, pH and buffers: insights from electrophoretic mobility and pH measurements of lysozyme solutions. *RSC Advances* **3**, 5882-5888 (2013).
35. Szymański, J.d. et al. Net charge and electrophoretic mobility of lysozyme charge ladders in solutions of nonionic surfactant. *The Journal of Physical Chemistry B* **111**, 5503-5510 (2007).
36. Yang, D., Kroe-Barrett, R., Singh, S. & Laue, T. IgG Charge. *Preprints* (2018).
